# Different time encoding strategies within the medial premotor areas of the primate

**DOI:** 10.1101/2023.01.28.526038

**Authors:** Hugo Merchant, Germán Mendoza, Oswaldo Pérez, Abraham Betancourt, Pamela Garcia-Saldivar, Luis Prado

## Abstract

The measurement of time in the subsecond scale is critical for many sophisticated behaviors, yet its neural underpinnings are largely unknown. Recent neurophysiological experiments from our laboratory have shown that the neural activity in the medial premotor areas (MPC) of macaques can represent different aspects of temporal processing. During interval categorization, we found that preSMA encodes a subjective category limit by reaching a peak of activity at a time that divides the set of test intervals into short and long. We also observed neural signals associated with the category selected by the subjects and the reward outcomes of the perceptual decision. On the other hand, we have studied the behavioral and neurophysiological basis of rhythmic timing. First, we have shown in different tapping tasks that macaques are able to produce predictively and accurately intervals that are cued by auditory or visual metronomes or when intervals are produced internally without sensory guidance. In addition, we found that the rhythmic timing mechanism in MPC is governed by different layers of neural clocks. Next, the instantaneous activity of single cells shows ramping activity that encode the elapsed or remaining time for a tapping movement. In addition, we found MPC neurons that build neural sequences, forming dynamic patterns of activation that flexibly cover all the produced interval depending on the tapping tempo. This rhythmic neural clock resets on every interval providing an internal representation of pulse. Furthermore, the MPC cells show mixed selectivity, encoding not only elapsed time, but also the tempo of the tapping and the serial order element in the rhythmic sequence. Hence, MPC can map different task parameters, including the passage of time, using different cell populations. Finally, the projection of the time varying activity of MPC hundreds of cells into a low dimensional state space showed circular neural trajectories whose geometry represent the internal pulse and the tapping tempo. Overall, these findings support the notion that MPC is part of the core timing mechanism for both single interval and rhythmic timing, using neural clocks with different encoding principles, probably to flexibly encode and mix the timing representation with other task parameters.

## Introduction

Time is a crucial parameter in life, and organisms have developed different mechanisms to quantify and predict events within the continuous flow of change in the environment (Tsao et al., 2022). Even if the central nervous system does not have a time sensory organ; animals are able to extract temporal information from stimuli of all sensory modalities and use it to generate timed behaviors. This paper focuses on the neural underpinnings of temporal processing during the perception and production of intervals in the hundreds of milliseconds range. This time scale is involved in basic but highly important behaviors observed since the invertebrates, such as collision avoidance and moving target interception (Merchant et al., 2001, 2003a, 2004a,b; Merchant & Georgopoulos, 2006). In addition, the range of hundreds of milliseconds is the scenario of complex behaviors such as the perception and production of speech (Assaneo et al., 2021), the execution and appreciation of music and dance (Merchant et al., 2015a; Mendoza & Merchant, 2014), and the performance of a large variety of sports (Merchant et al., 2003b). Researchers have found evidence of two distinct timing mechanisms in this scale: interval- and beat-based timing(Grube et al., 2010a,b; Teki et al., 2011; Breska & Ivry, 2018).In theory, interval-based timing implies the measurement of the absolute duration of discrete time intervals. In contrast, beat-based timing implies the quantification of relative durations with respect to the temporal regularity of the beat present in a stream of stimuli, such as a piece of music (Merchant et al., 2015a). Functional imaging and patient experiments have shown that the olivocerebellar and cortico-thalamic-basal ganglia circuit (CTBGc) are involved in interval and beat based timing, respectively (Teki et al., 2012; Grahn & Rowe, 2009; Sánchez-Moncada et al., 2020; Merchant & Bartolo, 2018; Cadena-Valencia et al., 2018). Nevertheless, the medial premotor areas (MPC), composed by the supplementary motor area proper (SMA) and the pre-supplementary motor area (preSMA), are the common output to both the cerebellar and basal ganglia circuits(Rajendran et al., 2018; Schwartze et al., 2012), conferring them the ability to process temporal information during both single interval and rhythmic perception and production tasks.

In this paper, we argue that neural populations in the primate MPC show flexible and multiplexed time encoding strategies that support both interval-based and beat-based timing, largely drawing from observations from our laboratory (see Merchant & Honing, 2014 for an initial view on this subject). In section 1, we describe the strategies by which preSMA encodes interval duration, category (long or short), and reward outcome based on a single-interval categorization task. In section 2, we show how neural populations in the MPC encode beat-based timing through a metronome synchronization-continuation task. We conclude with a brief summary and outlook for future work on timing neurophysiology in the primate brain.

### 1 Neurophysiology of Interval-based perception in MPC

Different psychophysical tasks have been developed to study how the brain processes temporal information from single intervals. We can classify these tasks according to their sensory and motor requirements or by the implicated sensory modality (Merchant et al., 2008c,b,a). In the motor domain, for example, one of the most employed paradigms is interval reproduction, in which subjects are first shown one interval or duration and then are asked to reproduce the duration with some motor response (Woodrow, 1930; Bartolo & Merchant, 2009; Jazayeri & Shadlen, 2015). Interval discrimination is one of the most employed paradigms in the sensory domain (Wearden, 1992; Kononowicz & van Rijn, 2014; Fabio et al., 2011). In this task, subjects are presented with two single, consecutive intervals and then are asked to emit a relative judgment about the two durations (Kim et al., 2013). The categorization of intervals as short or long is another instance of perceptual, interval-based timing (Ng et al., 2011; Méndez et al., 2014). This paradigm, also known as time bisection, is widely used since the late 70s of the past century to explore the mechanism by which animals perceive the passage of time (Church & Deluty, 1977; Gibbon, 1981; Allan & Gibbon, 1991; Wearden, 1991). In these tasks, subjects are initially trained to make one or another action in response to prototypic short or long intervals. When subjects learn to differentiate the short from the long intervals, intermediate test durations are presented. The subject’s goal is to categorize each test interval as short or long using the appropriate behavioral response. Despite being the subject of a long history of psychophysical studies, the neural basis of interval categorization remained unknown until recently.

#### 1.1 The ability of monkeys and humans to categorize intervals is similar

To provide information on the psychophysics and neural mechanisms of interval timing in primates, we developed a paradigm where human subjects and Rhesus monkeys (Macaca mulatta) categorize single intervals in the range of hundreds of milliseconds as short or long according to an imposed category limit (Méndez et al., 2011; Mendoza et al., 2018). In each task trial, the subjects were shown a test interval, with the onset and offset indicated by a brief visual stimulus displayed on a screen. After the interval offset and a fixed delay, the subjects communicated their perceptual decision by moving a cursor displayed on the screen into an orange circle, if the interval was categorized as short, or into a blue circle if the interval was classified as long. Crucially, subjects categorized three sets of eight intervals in every experiment. The short-long limit was the mean of the intervals of the set. Consequently, the shorter four intervals in a set should be categorized as short and the remaining as long. Importantly, the interval corresponding to the actual category limit of each set was never presented to the subjects as a test interval. As a result of this experimental design, the subjects had to change their subjective category boundary to classify the intervals of the different sets correctly.

We found that humans and monkeys perform similarly in these categorization tasks. Both species showed sigmoid psychometric functions, with the probability of long responses increasing as a function of the length of the intervals. In addition, both species got more correct answers for each block’s shortest and longest intervals and more decision errors for the intermediate intervals. Furthermore, for both humans and monkeys the bisection point, the interval at which the probability of ‘long’ response is 0.5, was close to the mean of the test durations of each set. Also, both species showed similar relative thresholds and an increase in temporal variability as a function of the implicit interval, following the Scalar Property for Timing (Gibbon, 1981; Allan & Gibbon, 1991). All these results were concordant with previous behavioral data from monkeys and humans categorizing time intervals and showed that both species have similar abilities for the categorical perception of time (Kopec & Brody, 2010; Merritt et al., 2010; Méndez et al., 2011). When human subjects were tested in our interval categorization task, we found that even though there was a 1-second-long delay between interval presentation and decision communication, categorization difficulty affected subjects’ performance, as well as their reaction and movement time. In addition, reaction and movement times were also influenced by the distance between the targets. This implies that not only perceptual, but also movement-related considerations were incorporated into the decision process Méndez et al. (2014). In addition, we conducted a beta burst TMS study in humans and found that decreasing the excitability of MPC produced a clear disruption of timing during the same categorization task (Méndez et al., 2017). Then we decided to study the neural properties of the primate preSMA cells during this paradigm.

#### 1.2 The subjective limit between the short and long categories is encoded by neurons of the monkey’s preSMA

We recorded the activity of neurons in the preSMA of Rhesus monkeys working in the relative interval categorization task. The monkeys had to change their subjective boundary within three blocks to correctly classify the intervals of the different sets. Interestingly, we found that this internal criterion is encoded by neurons in preSMA (Mendoza et al., 2018). We observed that some preSMA neurons, further called ‘boundary neurons’, generated a peak of activity at a relatively constant time after the interval onset, regardless of the actual duration of the presented interval. Thus, the peak of activity tended to occur after the interval offset when the test interval was short but before the interval offset when the test interval was long (Figure 1a). This timed activity could serve as a mental reference to delimit the duration of the short and long intervals. Such a mechanism would only require information of the relative time of occurrence of the peak activity and the interval offset. This hypothesis was supported by the fact that the time of occurrence of the peak activity changed according to the interval set being tested. Hence, the time at which neurons reached the peak activity was close to the actual category boundary of the current set. We observed this effect at the population and single neuron levels. For the population of boundary neurons, the mean of peak activity times was close to the actual boundary of each interval set. At the single neuron level, some neurons changed their time of peak activity across the different sets, occurring earlier for the shorter set and later for the longer set but always resembling the actual category limit (Figures 1a and 1b)

**Figure 1.**
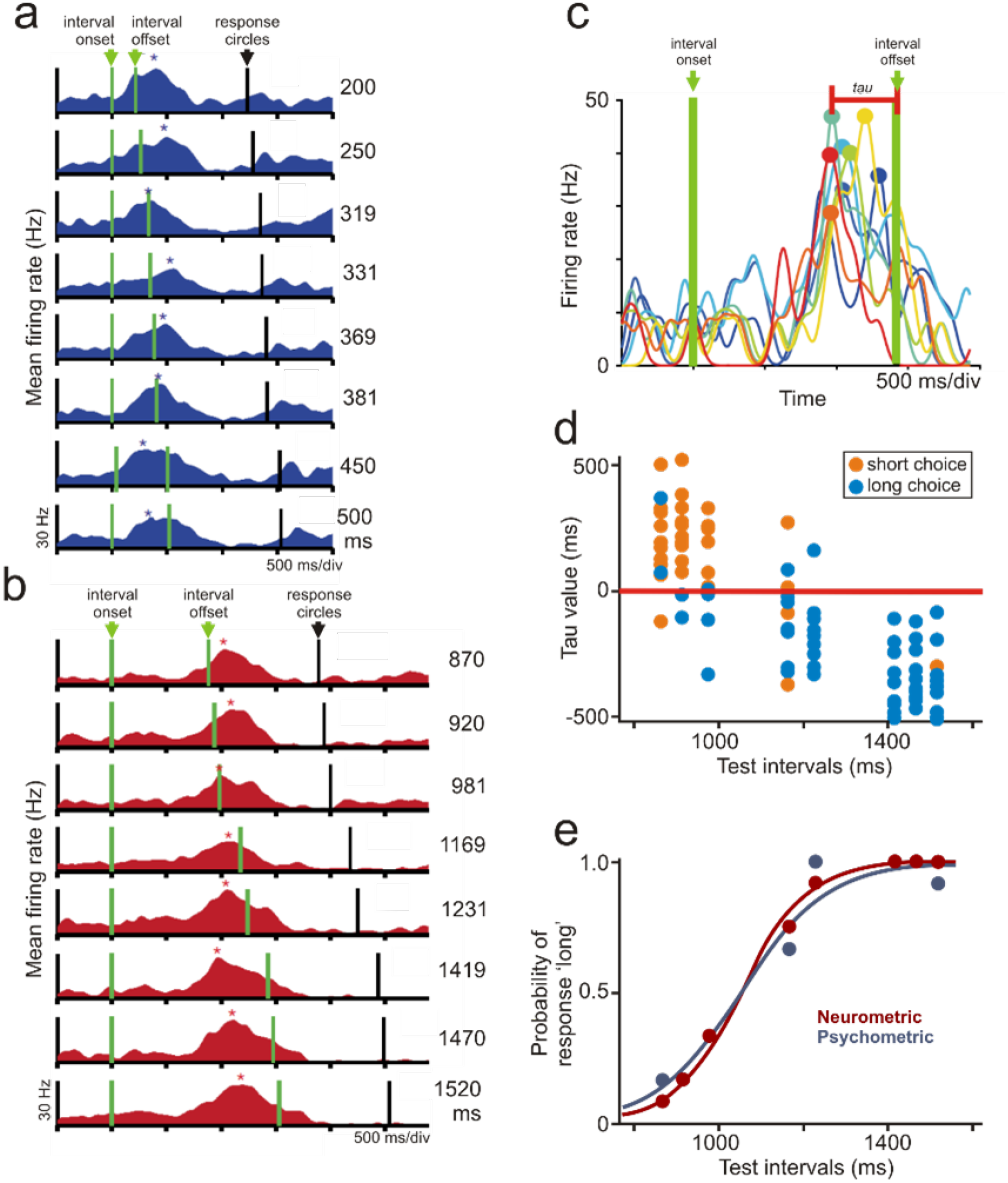
Neurons in the preSMA of the Rhesus monkey encode the main variables of the interval categorization task. a) Some preSMA neurons generate peaks of activity (indicated by the asterisks) after the offset of the short intervals but before the offset of the long intervals. Note the activity is segregated by the intervals being categorized (indicated to the right of the panels).b) Activity of the same neuron shown in a) but during the categorization of a different set of longer intervals. Note the peak activity occurs now around 1100 ms after interval onset. c) For each trial, the time difference (*τ*) between the peak of neuronal activity and the end of the test interval was measured. d) These measures tended to be positive for trials in which short intervals were presented and negative for long-interval trials and segregated short from long intervals in a way that correlated with the decision of the monkeys (inset). The red line indicates the best criterion for this boundary neuron and set of intervals. e) Psychometric and neurometric functions for the data shown in d). All figures adapted from Mendoza et al. (2018).

If boundary neurons encoded the subjective boundary, a correlation between the category selected by the monkeys and the category decoded from the activity of the boundary neurons should exist. Therefore, we determined whether the relative temporal occurrence of the peak activity and the end of the interval signaled by the presentation of the second stimulus can be used to decode the categorical decision of the monkeys. We quantified, for each boundary neuron and trial, the time elapsed (*τ*) between the peak of activity and the interval offset (Figure 1c). Next, we looked for the *τ* that minimized the error in classifying each interval as short or long (the best decoding criterion; Figure 1d). Finally, we constructed neurometric curves with the probability of a particular interval being categorized as long obtained from the proportion of trials in which *τ* was smaller than the best decoding criterion (Figure 1e). We found that the best criterions allowed the decoding of the monkey’s responses with high precision. Consequently, the relative thresholds and the points of subjective equality from the neurometric and the corresponding psychometric curves were similar. Significant trial-by-trial correlations between the category selected by the monkeys and the category decoded from the peak activity were found for a large group of neurons. This is remarkable since limit was never shown as a test interval to the monkeys. Therefore, the neural representation of the boundary was probably computed subjectively from the actually presented intervals.

#### 1.3 PreSMA neurons explicitly represent the categorical response of the monkeys

Since activity of boundary neurons can be used to decode the category selected by the monkeys, these results strongly support the notion that this signal encode a mental criterion to judge the intervals as short or long. Interestingly, we also found a sub-population of preSMA neurons that explicitly encode the category selected by the monkeys (Figure 1f). These category-encoding neurons showed selective activity to the short or long responses: their activity was similar for all the intervals assigned by the monkeys to the same category and different for the intervals classified in the opposite category. We found neurons with higher activity for intervals categorized as short and neurons with higher activity for intervals categorized as long.These responses are similar to previously reported neurons encoding the categorical decision in the frontal lobe and the basal ganglia during tactile (Romo et al., 1995, 1996; Merchant et al., 1997) and visual categorization tasks (Freedman et al., 2001; Merchant et al., 2011a). In contrast, no such signals are observed early in the sensory processing (Romo et al., 1996; Freedman et al., 2003).

#### 1.4 The reward outcome of the perceptual decisions is encoded by the preSMA

The consequences of the perceptual decision were also represented in the activity of preSMA neurons. In this case, some neurons showed activity selective to whether the monkeys’ responses were correct. This outcome-selective activity was observed after the monkeys’ responses, i.e., after the delivery, or absence, of reward. Hence, some outcome-encoding neurons were more active for correct responses, and other neurons showed higher activity for incorrect responses (Mendoza et al., 2018). This error signal may be essential in adjusting the subjective boundary or the choice during the learning or execution of the task. Further modeling studies are needed to reveal how the encoding of categorization variables emerges and is optimized in the preSMA neural circuits.

Overall, our results show a sequence of information encoding in preSMA during the categorization of intervals. First, the learned mental category boundary is evoked and used as a time reference to judge the ongoing interval.

Next, an explicit representation of the selected category emerges and is maintained as a memory trace during the delay previous to the actual response of the monkey. Finally, a strong representation of the outcome of the perceptual decision is generated and maintained during the inter-trial time (Figure 2c). Importantly, although the category boundary is encoded from the level of single neurons, the actual monkey’s criterion must result from the combined neural activity of all the population of boundary neurons. In fact, we found that the mean times of peak activity of all the boundary neurons correlated with the mean behavioral bisection point of the monkeys for each set of the categorization task (Mendoza et al., 2018). The concept of a population code may also apply to the neural representations of the selected category and the outcome of the perceptual decisions.

**Figure 2.**
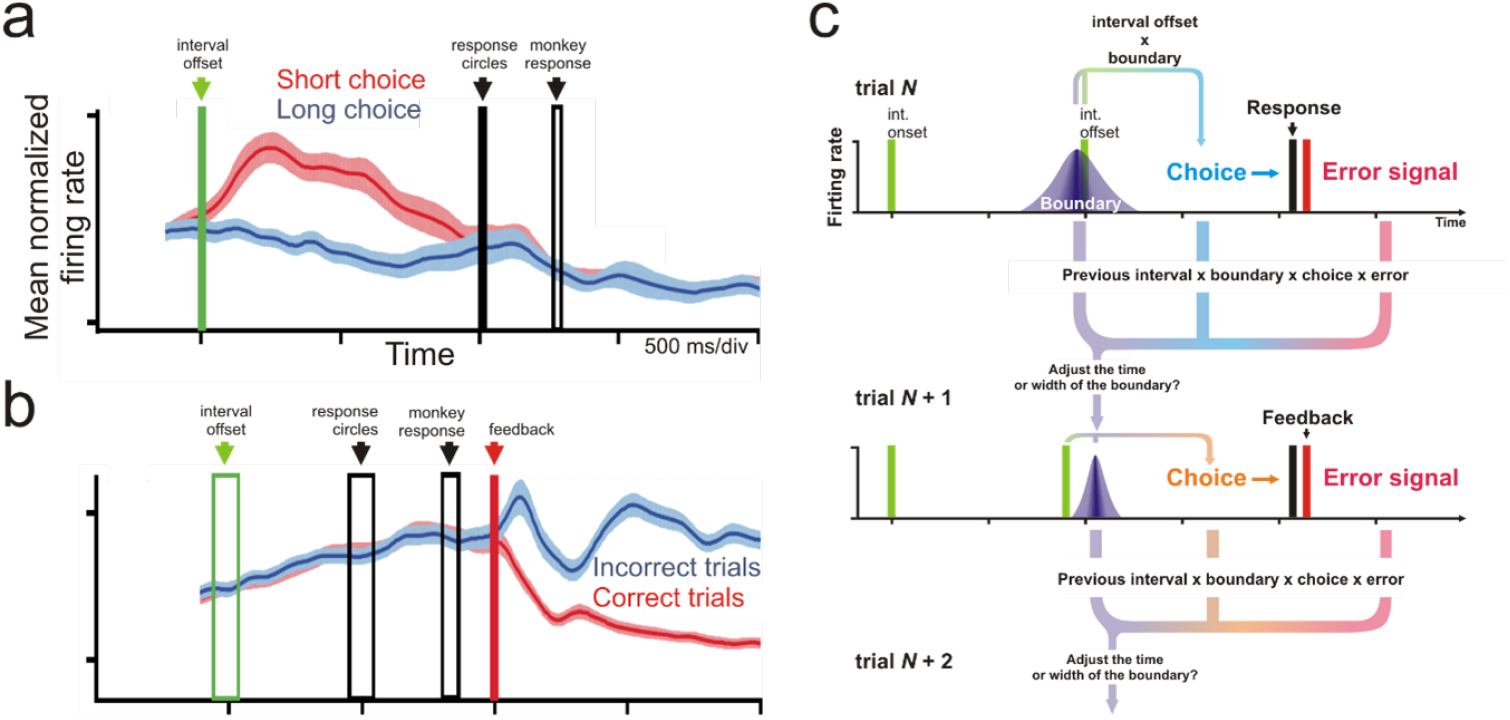
a) Mean (+ SEM) activity of a population of preSMA neurons encoding the category selected by the monkeys. In the short-response trials, these neurons showed higher activity before the actual monkey’s response. Note that the activity is aligned to the interval offset; the interval onset is not shown. b) Mean activity of a population of neurons encoding the outcome of the monkey’s decision. These neurons decreased their activity after the reward delivery (feedback time) in the correct trials. Activity is aligned to the time of feedback. c) Hypothetical trial-by-trial adjustment of the neural category boundary. In every trial of the task, intervals are judged to be short or long based on the subjective boundary (upper panel). After feedback, the time or width of the peak activity is adjusted depending on the task’s variables in the previous trials (lower panels). Figures a) and b) were adapted from Mendoza et al. (2018).

#### 1.5 Behavior and neurophysiology suggest interval categorization is solved by similar neural mechanisms in monkeys and humans

A key question is whether our experimental observations in the Rhesus monkey can be extrapolated to other animals, including humans. The concept of an internal criterion was proposed in 1981 to explain the behavior of different species in interval categorization (Gibbon, 1981). Subsequent behavioral and theoretical studies proposed that such a hypothetical criterion corresponds to the bisection point in humans and other animals and that is determined by the range of the test intervals (Killeen et al., 1997; Allan, 2002b). Nevertheless, the criterion hypothesis remained one of several possible alternative explanations (Gibbon, 1981; Maddox & Ashby, 1993; Allan, 2002a).Neurophysiologic studies in humans provided additional support to the idea of an internal criterion in humans. Lindbergh and Kieffaber Lindbergh & Kieffaber (2013) observed significant differences in the ERPs time-locked to probe offset between intervals judged to be short and long. In another study, Ng et al. (2011) reported an ERP, the Contingent Negative Variation, to increase in amplitude up to the value of the short prototype to remain at a constant level until about the mean of the short and long anchors and then, to return to baseline.These observations are consistent with a decision mechanism based on a category limit; i.e., during a trial of the categorization task, once the subjective time limit between the short and long intervals is exceeded, there is no need to continue attending the interval duration. Therefore, the category membership of the long intervals can be determined as early as the interval reaches the internal time criterion. This is consistent with the finding of shorter reaction times for the longer test durations in categorization tasks compared to other perceptual timing tasks, and with the observation that interval categorization is less demanding than other timing tasks that are supposed to require the tracking of the complete intervals (Bannier et al., 2019). Consequently, these behavioral observations, models, and neurophysiologic studies suggest that the neural mechanism operating in humans during interval categorization is similar to the mechanism we described in the monkey working in equivalent tasks.

#### 1.6 Factors influencing the subjective boundary

Experiments with human and non-human primates and other species performing interval categorization found choice biases resulting from manipulations of several task variables, including the range of the test intervals, the long-short ratio, the interval spacing and the probability of stimulus or reward occurrence (Elsmore, 1972; Stubbs, 1976; Allan, 2002b; Akdoğan & Balcı, 2016; Cambraia et al., 2019). Commonly, these manipulations produce systematic shifts of the psychometric functions with consequent changes in the bisection point. Notably, some of these variables were reported to produce similar changes in other perceptual tasks. In detection tasks, the psychometric thresholds are affected by manipulating the payoff or stimulus contingencies. This phenomenon, called the response bias, was proposed to depend on changes in the observer’s criterion (Gescheider, 1997).

A related phenomenon is the effect of the trial-by-trial history on the current subject’s performance. In interval categorization, the current judgments were reported to be sensitive to the choices in the previous trials (Wehrman et al., 2020).Thus, in order to learn the task or after changes in the task’s variables, the subjective limit must be fine-tuned by the events in the previous trials.

We suspect that the recent task history can bias the subject’s decisions and consequently can affect the peak activity of the boundary neurons. We propose that the time or width of the peak activity is modulated according to the variables in the previous trials (Figure 2c). With training, the peak activity of the boundary neurons would adapt to the statistics of the tasks, for example, to the range or probability of the test intervals or the reward contingencies. Therefore, adjusting the internal boundary would be an iterative process with critical importance during learning or after changes in task contingencies. An alternative explanation to the response bias or the ‘history effect’ is that they result from changes in the balance of the short-long choice-related neural activity, not from changes in the activity of boundary neurons. As we demonstrated, different neural codes in preSMA represent the subjective boundary and the selected category. Due to this functional segregation, our paradigm is well suited to determine whether specific task variables affect the internal criterion or the choice —ongoing studies in our laboratory attempt to provide data on these topics.

#### 1.7 No ramping activity in preSMA during perceptual timing tasks?

To end, we compare our observations during interval categorization with the neurophysiologic observations made in other timing tasks. Critically, we found that neurons in the monkey’s preSMA encode the main variables needed to solve the categorization task, but we did not find cells encoding elapsed time during the interval presentation. This is remarkable since different research groups have reported these types of activity in the preSMA of monkeys. Tanji and colleagues, reported that neurons in the SMA and preSMA of the Japanese monkey increased their activity as a function of the elapsed time (ramping activity) during the production of single intervals (Mita et al., 2009). Experiments in our laboratory with Rhesus monkeys trained in multiple-interval, synchronization-continuation tapping tasks also demonstrated ramping activity in preSMA during the interval production (Merchant et al., 2011b;see section 2 in this chapter). Interestingly, ramping activity related to elapsed time is also observed during the perceptual phase of interval reproduction tasks in other motor-related cortical areas. Each trial of these tasks has perceptual and reproduction phases. In the perceptual phase, a target interval is presented, and the subjects must attend to its duration but avoid movement. Then, in the subsequent reproduction phase of the trial, the subjects reproduce, with any motor action, the previously perceived interval(Jazayeri & Shadlen, 2015; Henke et al., 2021).

We hypothesize that ramping activity is mainly recruited by motor-related areas during tasks requiring the quantification of the whole intervals, such as the perceptual or motor phases of motor timing tasks or perceptual discrimination tasks (references). Not only does interval categorization lack an interval reproduction phase, but our decoding analysis also demonstrated that it could be solved without quantifying the total interval duration. These ideas are consistent with most neurophysiologic studies in monkeys (Perrett, 1998; Maimon & Assad, 2006; Renoult et al., 2006; Tanaka, 2007; Lebedev et al., 2008; Genovesio et al., 2009; Jazayeri & Shadlen, 2015; Mendoza et al., 2018; see also Oshio et al., 2008). Nevertheless, we cannot discard the possibility that elapsed time can be decoded from the activity of neurons in other brain areas (see for example Gouvêa et al., 2015 for the neurophysiology of interval categorization in the striatum of rats). Additional studies in different animal species, with a more diverse battery of timing paradigms and recording the simultaneous activity of several cortical areas, are needed to clarify these issues.

In conclusion, our data suggest that the monkey preSMA solves interval categorization using a simple mechanism that does not require exhaustive quantifying each interval but fine-tuning a time boundary. Initial evidence from different research groups suggests that the same mechanism might operate in the human brain. Therefore, our observations support classical psychophysical hypotheses suggesting that subjects compare the intervals to be categorized with an internal criterion that depends on the distributions of the test intervals, the choice biases, and the reward contingencies.

### 2 Neurophysiology of Beat-based perception in MPC

#### 2.1 Psychophysics of rhythmic tapping

Music and dance depend on intricate loops of perception and action, where temporal processing can be engaged during the synchronization of movements with sensory information or during the internal generation of movement sequences (Janata & Grafton, 2003). Thus, beat induction is the cognitive ability that allows humans to hear a regular pulse in music and to move in synchrony with it. Importantly, without beat induction there is no music perception and, hence, is considered a universal human trait (Honing, 2012; Honing et al., 2012, 2018). These cognitive abilities depend on an internal brain representation of pulse that involves the generation of regular temporal expectations, which is the core of the beat-based timing mechanism (Balasubramaniam et al., 2021). Several studies have shown that the internal pulse is directly mapped to the timing of the entrained movements, typically measured as tapping movements that occur few milliseconds before the beat (Lenc et al., 2021; Nozaradan et al., 2016, 2017). A classical task used to study beat induction is the synchronization-continuation tapping task (SCT), which can be considered a simplified version of beat perception and entrainment in music. In the SCT, subjects tap in sync with periodic sensory cues (synchronization epoch), and then keep tapping after the metronome is extinguished using an internal beat representation (continuation epoch) (Wing, 2002) Performance in this task shows two features present in other timing tasks: a linear increase in variability of produced intervals with the mean that is a form of Weber’s law, called scalar property (Gibbon et al., 1997; Merchant et al., 2008c; Garcia-Saldivar et al., 2021), aand an over- and underestimation of intervals for shorter and longer intervals, termed regression towards the mean or bias effect (Woodrow, 1934; McAuley & Jones, 2003; Pérez & Merchant, 2018). Furthermore, subjects use an error correction mechanism that maintains tap synchronization with the metronome, since a longer produced interval tends to be followed by a shorter interval and vice versa, to avoid error accumulation and losing the metronome (Repp, 2005; Iversen et al., 2015; Perez et al., 2023 in preparation). In contrast, during continuation, there is a drift in the duration of produced intervals (Madison, 2001; Collier & Ogden, 2004).

Humans possess a remarkable flexibility for beat-based timing, recognizing the beat from a wide range of complex rhythms, and with the natural tendency to predictively entrain to the beat by moving different body parts, such as finger or foot taps (Patel, 2018).Recently, it has been demonstrated that macaques possess the neural machinery to perceive and entrain to the simplest form of beat: an isochronous auditory metronome. On one side, beat perception has been measured with mismatch negativity (MMN), an auditory event-related EEG potential that can be used as an index of a violation of temporal expectation. Notably, MMN is sensitive to violations of the beat using complex or simple rhythms in humans but only for isochronous metronomes in monkeys (Honing, 2012; Honing et al., 2018). On the other, psychophysical experiments showed that, when immediate feedback about the timing of each movement is provided, monkeys can predictively entrain to an isochronous beat, generating tapping movements in anticipation of the metronome (Gámez et al., 2018; see also García-Garibay et al., 2016; Takeya et al., 2017). In fact, macaques can flexibly change their movement tempo from trial to trial covering a range from 400 to 1000 ms (unpublished observations). Furthermore, monkeys can superimpose accentuation patterns onto an isochronous auditory sequence, suggesting that they can generate a simple subjective rhythm on a regular auditory sequence (Ayala et al., 2017; Criscuolo et al., 2021). Thus, both primate species can produce negative asynchronies, show an error correction mechanism, produce precise and accurate interval during synchronization to an isochronous (Betancourt et al., 2022). Nevertheless, humans perform better with auditory but monkeys with visual metronomes (Zarco et al., 2009; Gámez et al., 2018; Betancourt et al., 2022). These findings support the gradual audio-motor hypothesis that suggests that beat-based timing emerged gradually in primates, peaking in humans due to a sophisticated audio-motor circuit, and that is also present for isochrony in macaques where it depends on the close interaction between MPC, CTBGc, and the auditory cortex (Honing & Merchant, 2014; Merchant & Honing, 2014). Indeed, in both primate species, it has been shown that MPC plays a critical role in beat extraction and entrainment (Rao et al., 1997; Chen et al., 2008; Merchant et al., 2015a; Mendoza & Merchant, 2014).

#### 2.2 Ramping activity as a single cell timing signal for rhythmic tapping

We recorded the activity of MPC cells during a version of the SCT where monkeys produced three intervals in the synchronization and three intervals in the continuation epochs (Figure 3a). Brief auditory or visual interval markers were used during the synchronization phase and the range of target intervals was from 450 to 1000 ms (Zarco et al., 2009). The monkeys were able to produce the target intervals accurately, showing an average underestimation of 50 ms across interval durations during the synchronization and continuation phases of the SCT. In addition, we analyzed the temporal variability of the monkeys’ tapping performance, defined as the SD of the individual inter-response intervals (Merchant et al., 2008c,b). The temporal variability followed the scalar property, with a linear increase as a function of interval duration in both phases of SCT (Zarco et al., 2009). Furthermore, the analysis of the tapping hand kinematics revealed that monkeys temporalize the pauses or dwell between movements, while producing stereotypic downward and upward movements with a similar duration across the tested metronome tempos (450-1000 ms). These findings suggest that monkeys use an explicit timing strategy to perform the SCT, where the timing mechanism controls the dwell duration, while also triggering the execution of stereotyped tapping movements across each produced interval in the rhythmic sequence (Donnet et al., 2014; Gámez et al., 2018).

**Figure 3.**
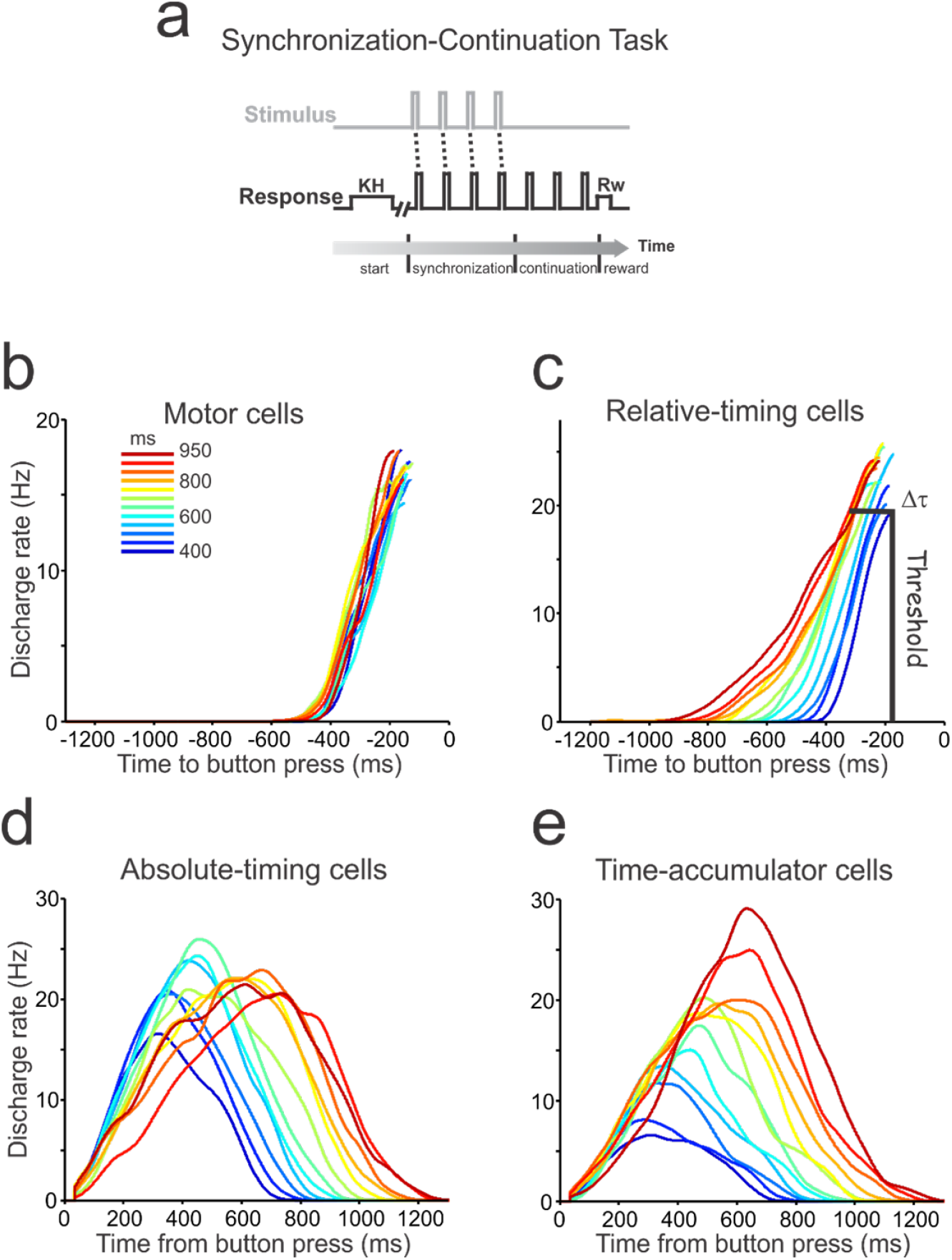
(a) Synchronization-Continuation Task (SCT). Monkeys were required to push a button (R, black line) each time stimuli with a constant interstimulus interval (S, gray line) were presented, which resulted in a stimulus-movement cycle. After four consecutive synchronized movements, the stimuli stopped, and the monkeys continued tapping with similar pacing for three additional intervals. The target intervals, defined by brief auditory or visual stimuli, were 450, 550, 650, 850, and 1,000 ms, and were chosen pseudo-randomly within a repetition. Different ramp populations: motor (b), relative-timing (c), absolute-timing (d), and time-accumulator (e) cells. b and c are aligned to the next button press, while c and d are aligned to the previous button press. The duration of produced intervals is color-coded in b. All these ramp population functions correspond to the addition of individual ramps over time. Modified from (Merchant et al., 2011b).

The extracellular activity of single MPC neurons was recorded during task performance. A large population of neurons showed ramping activity before or after each sensory or motor event in the SCT (Merchant et al., 2011b). Consequently, we developed a warping algorithm to determine whether the responses of the cells were aligned to the sensory or motor aspects of the SCT, and we found that most MPC cells were aligned to the tapping movements instead of the stimuli used to drive the temporal behavior (Pérez et al., 2013; Merchant et al., 2015b).

Next, we used an iterative algorithm to find the best regression model to explain the increase or decrease of instantaneous activity over time with respect to the tapping times using the spike density function. With this method, we defined for each ramp the following parameters: duration, slope, peak magnitude, and the time τ from the peak to the button press. Using this information, we classified different cell populations with ramping activity into four groups: motor, relative-timing, absolute-timing and time-accumulator, and swinging cells (Merchant et al., 2011b). A large group of cells shows ramps before the movement onset that are similar across produced durations and the sequential structure of the task and, therefore, are considered motor ramps (Figure 3b). Interestingly, another cell population showed an increase in ramp duration but a decrease in slope as a function of the animals’ produced duration, reaching a similar discharge magnitude at a specific time before the button press. These cells are called relative-timing cells since their ramping profile could signal how much time is left to trigger the button press in the task sequence (Figure 3c). Therefore, this population of MPC neurons has the response properties to encode the time remaining for an event or time-to-contact cells, and once the population reaches a firing magnitude threshold, it could trigger the internal beat signal (Merchant et al., 2011b).

Other groups of cells show a consistent increase followed by a decrease in their instantaneous discharge rate when their activity was aligned to the previous button press rather than to the next one. These cells showed an up-down activation profile whose duration increased as a function of the produced interval (Figure 3d) and were called absolute-timing cells. In addition, we found cells that responded as sand clocks since their activity was accumulated as a function of the passage of time, with final peaks of activity that increased linearly with the produced interval and hence were denominated time-accumulator cells (Figure 3e). Therefore, these cells could be representing the elapsed time since the previous movement, using two different encoding strategies: one functioning as an accumulator of elapsed time where the peak magnitude and the duration of the activation period are directly associated with the time passed, and another where only the duration of the activation period is encoding the length of the time passed since the previous movement (Merchant et al., 2011b; Merchant & Averbeck, 2017).

Cell activity changes associated with temporal information processing in behaving monkeys have been reported in the cerebellum (Perrett, 1998; Ohmae et al., 2017; Okada et al., 2022), the putamen (Bartolo et al., 2014; Bartolo & Merchant, 2015; Merchant & Bartolo, 2018) (Bartolo et al., 2014; Bartolo and Merchant, 2015; Merchant and Bartolo, 2018), the caudate (Kameda et al., 2019; Kunimatsu et al., 2018), the thalamus (Tanaka, 2007; Wang et al., 2018), the posterior parietal cortex (Leon & Shadlen, 2003; Maimon & Assad, 2006; Jazayeri & Shadlen, 2015), and the prefrontal cortex (Oshio et al., 2008; Brody, 2003; Henke et al., 2021), as well as in the motor cortex (Merchant et al., 2004b; Lebedev et al., 2008; Renoult et al., 2006), and MPC (Mita et al., 2009; Merchant et al., 2014; Sohn et al., 2019). These areas form different circuits linked to sensorimotor processing using the skeletomotor or oculomotor effector systems. Most of these studies have described climbing activity during different timing contexts, which include discrimination of time, time estimation, time categorization, single interval reproduction, and rhythmically produced saccades and hand movements. Therefore, the increase or decrease in instantaneous activity as a function of the passage of time is a property present in many cortical and subcortical areas of the CTBGc and the cerebellum that may be involved in different aspects of temporal processing in the hundreds of milliseconds scale. In fact, the ubiquitous presence of cells’ increases or decreases in discharge rate as a function of time across different timing tasks and areas of a potential core timing circuit suggests that ramping activity is a fundamental element of the timing mechanism.

The recently reported temporal scaling as a mechanism to encode time remaining for action is very evident in ramping cells that behave as time-to-contact cells or the absolute timing cells reported by us (3c,d Merchant & Averbeck, 2017). These cells show a similar instantaneous pattern of activations that is contracted for short and expanded for long intervals and can be the single-cell primordium for the neural population temporal scaling observed in neural trajectories in state space (Wang et al., 2018; Remington et al., 2018). In contrast, the time-accumulator cells. This type of cells might be very common during tasks that demand elapsed time encoding instead of predicting a sensory, cognitive, or motor event (Bi & Zhou, 2020; Merchant & Pérez, 2020).

#### 2.3 Interval tuning: a circuit signal for context dependent flexibility

Psychophysical studies on learning and generalization of time intervals support the notion that neurons in the timing circuit are tuned to specific interval durations but can be activated in a modality- and context-independent fashion (Meegan et al., 2000; Nagarajan et al., 1998; Bartolo & Merchant, 2009; Sánchez-Moncada et al., 2020). Accordingly, we found a graded modulation in the discharge rate of cells as a function of interval duration during the SCT in cells of MPC (Merchant et al., 2013a). Figure 4a,b shows the profile of activation of a cell in the preSMA of a monkey performing this task. The neuron shows larger activity for the longest durations, with a preferred interval of around 900 ms. A large population of MPC cells is tuned to different interval durations during the SCT, with a distribution of preferred intervals that covers all the tested durations, although there was a bias towards long preferred intervals (Merchant et al., 2013b). These observations suggest that the MPC represents interval duration, where different populations of interval-tuned cells are activated depending on the duration of the produced interval (Merchant & Bartolo, 2018). In addition, most of these cells also showed selectivity to the sequential organization of the task, a previously described in sequential motor tasks in MPC (Tanji, 2001). The cell in Figure 4a,b also shows an increase in activity during the fifth produced interval in the SCT sequence.

**Figure 4.**
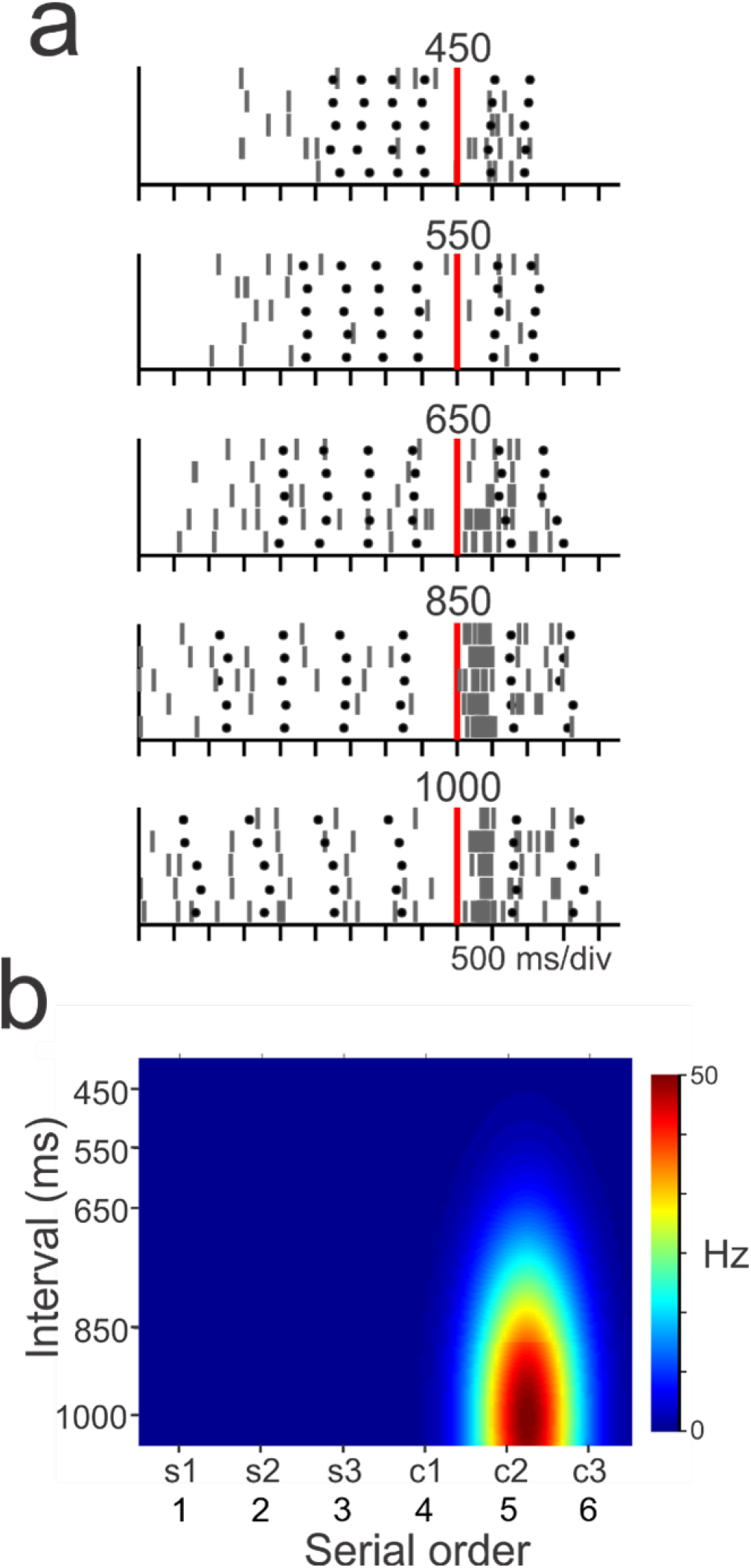
(a) Responses of a sharply double-tuned MPC cell with a long-preferred interval and a preferred sequence order around the second continuation interval. Circles correspond to tap times. The raster is aligned (red line) to the second tap of the continuation phase. All target intervals are showed in a vertical arrangement. (b) Double-Gaussian tuning function for the cell responses depicted in a. The heatmap discharge scaler is on the right. Modified from (Merchant et al., 2013a).

Again, at the cell population level, all the possible preferred ordinal sequences were covered (Merchant et al., 2013b). Hence, the temporal and sequential information is multiplexed in a cell population signal that defines the duration of the produced interval and its position in the learned SCT sequence (Merchant et al., 2013a; Bartolo et al., 2014). Overall, these findings support the notion that MPC uses mixed selectivity to represent the passage of time, the tempo duration, and the serial order during SCT (Merchant et al., 2013b; Merchant & Bartolo, 2018; Gámez et al., 2019).

Interval tuning during single interval and beat based timing has been reported in MPC (Merchant et al., 2013b; Mita et al., 2009), the prefrontal cortex (Henke et al., 2021), the putamen (Bartolo et al., 2014; Bartolo & Merchant, 2015), the caudate (Kameda et al., 2019; Kunimatsu et al., 2018), and the cerebellum (Ohmae et al., 2017; Okada et al., 2022). In addition, a chronomap in the medial premotor cortex has been described in humans using functional imaging. The interval-specific circuits show a topography with short preferred intervals in the anterior and long preferred intervals in the posterior portion of SMA/preSMA (Protopapa et al., 2019). Hence, timing not only depends on one population of cells that contracts or expands their activity patterns depending on a constant speed knob (Wang et al., 2018) but also on interval-specific neurons that build distinct timing circuits. It is well known that tuning and modularity are mechanisms for the division of labor that are used in cortical and subcortical circuits to represent sensory, cognitive and motor information (Hubel & Wiesel, 1977; Vernon B, 1998; Georgopoulos et al., 2007; Goldman-Rakic et al., 1984; Naselaris et al., 2006a,b). Consequently, interval tuning can provide large flexibility to mix time encoding and prediction with other task parameters, which are also represented in the premotor system but with different mapping frameworks (Merchant & Yarrow, 2016; Merchant & Bartolo, 2018; Zhou et al., 2022).

#### 2.4 Neural sequences

As a population, ramping and/or tuned MPC cells show activation profiles that are far from static. Indeed, these cells are recruited in rapid succession producing a progressive neural pattern of activation (called neural sequences or moving bumps) that flexibly fills the beat duration depending on the tapping tempo. Hence, neural sequences provide a relative representation of how far an interval has evolved between the taps (Crowe et al., 2014; Merchant et al., 2015b; Zhou et al., 2020). Notably, this periodic clock resets on every period cycle, encoding the interval pulse in this resetting. In fact, the neural chain progression starts with a group of cells, migrates to other cells during the timed interval, stops with the last group of cells, and simultaneously is initialized for the next produced interval with the previous initial set of cells (Merchant et al., 2015a; Gámez et al., 2018; Lenc et al., 2021). Three parameters of the moving bumps are directly linked with the representation of the tapping tempo duration: (1) The duration of the activation period for each cell within the moving bump, (2) the number of neurons comprising the neural sequence, and (3) neural recruitment lapse, which is the time between pairs of consecutively activated cells (see Figure 5c). We have shown that the rhythmic neural clock in MPC uses a mixed encoding strategy between temporal scaling or absolute time encoding to represent the tempo. Under the temporal scaling scenario, the activation profile of a neuron in the moving bump is the same between target durations but shrinks for short and elongates for longer tempos. Under the absolute timing strategy, the activation periods are the same across durations, but additional neurons are recruited for longer durations so that the new neurons are active in the last portion of the interval (Zhou et al., 2022). Our results indicate that both the duration and the number of neurons increased as a function of the target tempo in the task, indicating the presence of both types of encoding strategies (Merchant et al., 2015b; Gámez et al., 2019).

Neural sequences have been reported in the striatum (Mello et al., 2015; Gouvêa et al., 2015; Kim et al., 2013; Zhou et al., 2020; Bakhurin et al., 2017; Jin et al., 2009), prefrontal cortex (Kim et al., 2013; Tiganj et al., 2017) and medial premotor areas (Crowe et al., 2014; Merchant et al., 2015b; Gámez et al., 2019; Murakami et al., 2014) in perception and production interval- and beat-based timing tasks across primate and rodent species. Hence, this is a population code that is widespread across the CTBGc core timing network and the prefrontal cortex during different timing paradigms and maybe the most reliable neural population clock. In fact, a recent empirical and computational study suggested that neural sequences are an effective representation of time from the downstream readout point of view (Zhou et al., 2020).

**Figure 5.**
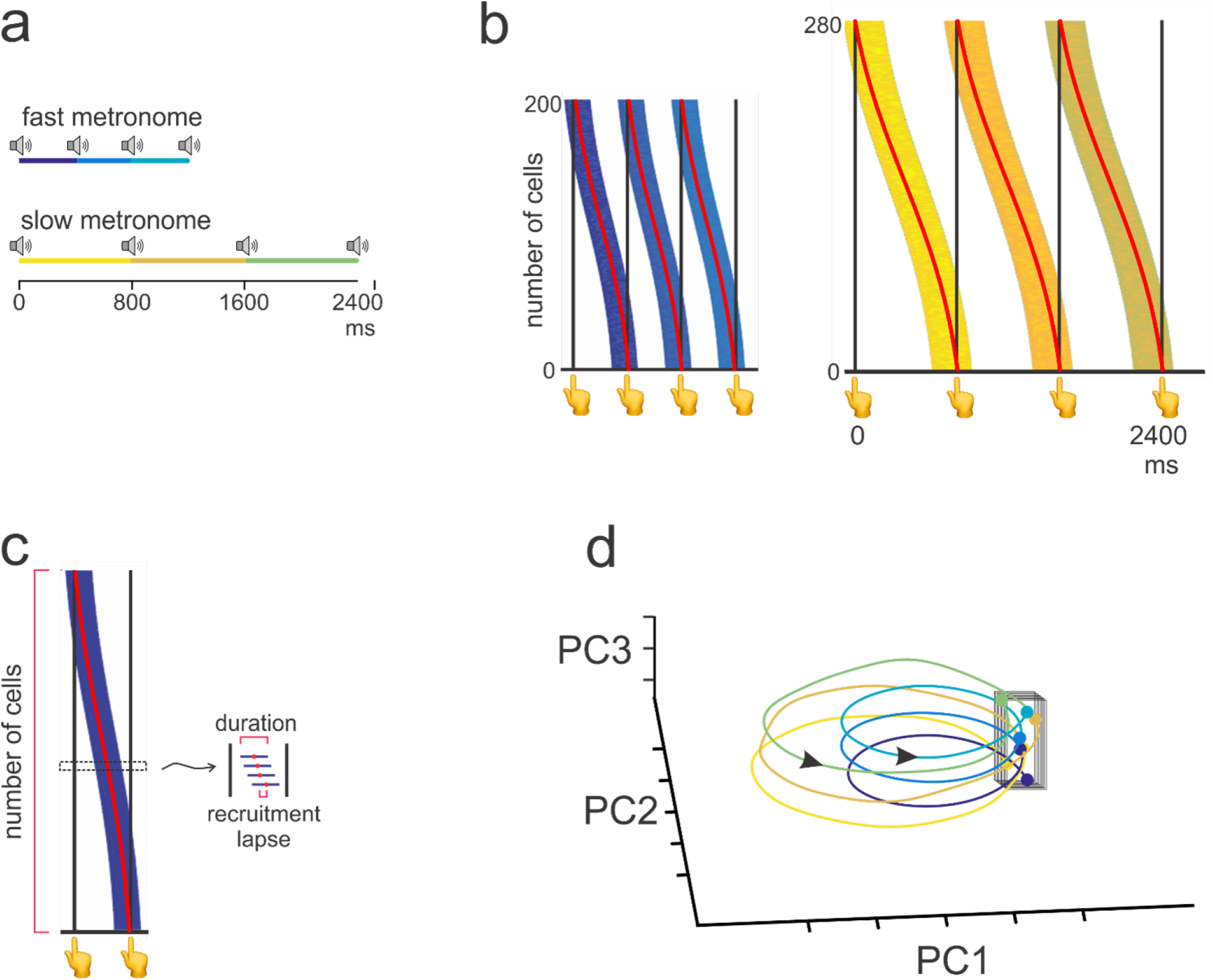
(a) Simplified version of the SCT with only three produced intervals in the synchronization epoch. The small speakers correspond to the isochronous stimuli for the fast and slow tempos. The color code of the three produced intervals is used in b and c to signal the timing neural population codes in the task sequence. (b) Neural sequences for the fast (top) and slow (bottom) tempos producing three regenerating moving bumps for each produced interval. Individually colored stripes are constituted by multiple horizontal lines, where each line corresponds to the onset and duration of the activation period for one cell. For the fast tempo we simulated 200 cells with a mean activation duration of 200 ms, while for the slow tempo we simulated 280 cells with a mean activation duration of 300 ms. All cells were sorted by their peak activation time. (c) The three key parameters of neural sequences are outlined in pink: the number of cells in the moving bump, the duration of the activation period of each neuron, and the recruitment lapse that corresponds to the time between the activation peaks of two consecutive neurons. (d) Neural trajectories during the synchronization task. The trajectory starts from a tapping separatrix (black cuboid), completes a cycle during every intertap interval, and returns to the tapping separatrix. The separatrix is invariant across durations and serial order elements of the task. The metronome’s tempo modulates the amplitude of the trajectories and the serial order element as the third axes in the state population, generating for each produced interval an evolving population pattern of activation. The circles correspond to the tapping times and the arrows specify the direction of the trajectories’ movement. Note that a similar population response profile is repeated cyclically for the three intervals (color coded) and that the resetting of each moving bump corresponds to a potential internal pulse representation.

#### 2.5 Neural population trajectories

The time-varying activity of cell populations can be projected into a low-dimensional state space using dimensional reduction techniques These techniques capture the covariance of the activity of the cell population across time and can reveal emergent properties that are not present in the single-cell activity (Elsayed & Cunningham, 2017). Indeed, the neural trajectories that we obtained using Principal Component Analysis (PCA) on more than a thousand MPC neurons recorded in the SCT show the following properties (Figure 5a,d) (Mendoza et al., 2016). First, they have circular dynamics that form a regenerating loop for every produced interval. Notably, the population state during pulse-based predictive timing correlates with the traversed proportion of an interval (relative timing) instead of its absolute magnitude (Gámez et al., 2019; Balasubramaniam et al., 2021). Second, the periodic trajectories increase in amplitude as a function of the tapping interval. These period-dependent increments in the trajectory radius result from a larger number of neurons within a moving bump (Gámez et al., 2019). Finally, the neural population trajectories converge in similar state space at tapping times, resetting the pulse-based clock at this point (Gámez et al., 2019) (Figure 5c). Hence, the convergence to this neural attractor state could be the internal representation of the pulse transmitted as a phasic top-down predictive signal to the auditory areas before each tap (Lenc et al., 2021).

The notion that beat-based timing depends on a population clock whose dynamics at each instant correspond to a unique pattern in state-space started 2014 (Merchant et al., 2014). In that article we suggested that on top of the single-cell encoding of time there was a population timer that represented the relative passage of time, the duration of the produced intervals, and the serial order elements of the tapping sequence during SCT. Since then, there have been many reports of neural population clocks during interval-based timing using neural trajectories in medial premotor areas(Wang et al., 2018; Remington et al., 2018; Sohn et al., 2019; Gámez et al., 2019), prefrontal cortex (Kim et al., 2013; Xu et al., 2014; Henke et al., 2021), and the striatum (Zhou et al., 2020). These neural clocks can compute elapsed time in the final position, time remaining for an action in the time scaling and, flexibly incorporate the task context into the kinematics of the neural trajectories (Remington et al., 2018; Egger et al., 2019; Henke et al., 2021; Lafuente et al., 2022). Therefore, the neural trajectory clock is now the most accepted neurophysiological mechanism to quantify and predict events in time in both interval- and beat-based timing tasks (Tsao et al., 2022). The parallelisms between the geometric and kinematic properties of neural trajectories and the properties of neural sequences have also been thoroughly documented (Betancourt et al., 2022).

#### 2.6 Integrating layers of neuronal clocks in the medial premotor cortex

Our neurophysiological recordings in behaving animals indicate that MPC, an area of the core timing mechanism (Merchant et al., 2013a), uses multiple encoding strategies and different organization levels to represent diverse aspects of the temporal and sequential structure of the SCT. The different types of ramping activity; the cells with mixed selectivity to tempo duration, serial order, and elapsed time; the progressive patterns of neuronal activation; and the neural trajectories must be interlinked to generate a coordinated population clock that flexibly processes temporal information across a wide variety of tasks. Consequently, it is crucial to generate metrics that define the rules of interaction between these temporal signals and to determine the bidirectional effects of lower to upper levels of the neural organization on timing. It is also fundamental to determine the interaction between the neural clock and other task components such as decision making, reinforcement learning, the calculation of value, and the impact of previous trials of the task-solving strategy. Important attempts to tackle the first issue come from our lab, where we have documented that different types of ramping activity are active at different stages of the neural sequences, that duration-tuned cells are mainly engaged in the intermediate part of the moving bumps, and that the geometry and kinematics of neural trajectories have a counterpart in the properties of the neural sequences (Gámez et al., 2019; Betancourt et al., 2022). Recently, the group of Thurley also linked the properties of mixed selectivity, ramping, neural sequences, and neural trajectories (Henke et al., 2021) during single interval measurement and reproduction in the prefrontal cortex of gerbils. Nevertheless, an integrated analysis on the timing mechanisms across tasks and species is lacking and urgently needed.

#### 2.7 Structural bases for beat-based timing during SCT

Although rhythmic entrainment is prevalent across all human cultures (Nettle, 2009), there are wide individual differences in the precision, accuracy, and predictability of movement synchronization (Garcia-Saldivar et al., 2021). A critical unanswered question is what the structural bases for these differences in SCT are. To address this issue, we first obtained diffusion-weighted images from human subjects who also performed the SCT with auditory or visual metronomes and five tempos ranging from 550 to 950 ms. Then, we developed a method to determine the fiber density of U-fibers running tangentially to the cortex (Garcia-Saldivar et al., 2021), which are the core fiber system for corticocortical associations (Schüz & Braitenberg, 2002). Notably, the right audio-motor system (including MPC) showed individual differences in the density of U-fibers that were highly correlated with the degree of predictive entrainment across subjects, measured with the asynchronies during the SCT. These correlations were selective for the synchronization epoch and were specific for auditory metronomes with tempos around 700 ms. In addition, we found that the predictive rhythmic entrainment abilities of subjects were significantly associated with the density and bundle diameter of the corpus callosum (CC), forming a chronotopic map where behavioral correlations of short and long intervals were found with the anterior and posterior portions of the CC (Garcia-Saldivar et al., 2021). Consequently, these findings support the notion that the structural properties of the superficial and deep white matter of the audio-motor system define the predictive abilities of subjects during rhythmic tapping.

A similar methodology can be applied to determine the structural changes in the white matter and the plastic modifications of cortical thickness due to intensive learning in non-human primates (Garcia-Saldivar et al., 2021; Messinger et al., 2021) (Poirier et al., 2021). Indeed, a longitudinal approach can be used to collect and analyze image data at different stages of learning to determine the effects of training on rhythmic perception and entrainment tasks in macaques (Milham et al., 2020; Song et al., 2021). Preliminary observations from our lab suggest that the density of superficial white matter in the cortical areas of the core timing network (Merchant et al., 2013a) undergoes learning-induced changes. Crucially, this structural information can be used in conjunction with behavioral and neurophysiological information to generate a multimodal map that combines all these data in the same structural framework. This integrated construct can be used to understand more deeply the anatomofunctional correlates of rhythmic behavior for each animal and across animals (Garcia-Saldivar et al., 2021).

### 3 Some considerations on the brain mechanisms behind interval- and beat-based timing

The recent myriad of neurophysiological studies on the neural basis of timing have mainly focused on the timing signals of the striatum and medial prefrontal cortex in rodents performing single interval tasks (Gouvêa et al., 2015; Tiganj et al., 2017; Mello et al., 2015; Zhou et al., 2020; Bakhurin et al., 2017; Jin et al., 2009). These papers have documented ramping activity, neural sequences, and trajectories with time scaling properties. The main principle behind these studies is that time can be represented as stable, repeatable activity patterns in single cells so that the overall population state activity systematically changes as a function of time within these two connected brain areas. This rule is followed on tasks in the range of hundreds of milliseconds and seconds(Tsao et al., 2022). On the other hand, the human imaging and lesion literature support the hypothesis of functionally non-overlapping mechanisms of interval and beat-based timing, with the separable contributions of the cerebellum and basal ganglia to these two types of temporal processing (Grube et al., 2010a; Teki et al., 2012; Breska & Ivry, 2018). Obviously, the rodent neurophysiology on single interval tasks in the striatum contradicts this hypothesis. All these studies claim that the basal ganglia are a key component of single interval timing. This discrepancy could be due to several factors, including: (1) the limited generalization in the brain anatomy of the rodent and the human, especially regarding the frontal lobe and the basal ganglia (Mendoza & Merchant, 2014); (2) the limited range of timing behaviors than can be trained in rodents, compared with the immense flexibility and behavioral repertoires in humans for measuring time; (3) the methodological restrictions of measuring the neural activity in human using the slow BOLD signal (seconds resolution) or the inherent problems of using behavior in neurological patients to understand brain mechanisms (Merchant et al., 2008a). Needless to say, the neurophysiology in non-human primates can partially address these issues since the anatomy, physiology, and behavioral spectrum in monkeys is quite close to that of humans (Merchant et al., 2003b, 2004c; Merchant & Honing, 2014; Mendoza & Merchant, 2014). In particular, invasive high-density electrodes can be placed in many areas simultaneously during task performance (Mendoza et al., 2016), obtaining neurophysiological signals that can reveal a potential neural clock generalizable to the Homo sapiens. Recordings in the primate putamen on single interval reproduction tasks have revealed neural signals with the same properties observed in rodents (Wang et al., 2018; Sohn et al., 2019); while experiments in monkeys performing a rhythmic event detection task have shown that the cerebellum provides more accurate time predicting information than the caudate (Kameda et al., 2019). Furthermore, in a rhythmic saccadic task where monkeys show predictive timing behavior (Takeya et al., 2017), the cerebellum shows a complex set of responses, including predictive rhythmic activity (Okada et al., 2022). Therefore, these studies clearly contradict the notion of a dissociation in the mechanisms for interval and beat based timing, and instead suggest that both the basal ganglia and the cerebellum play important roles in timing both single and periodic events in coordination with the medial premotor areas. Further experiments with simultaneous recordings across these circuits in macaques performing both single and rhythmic timing tasks are urgently needed to have a clear notion of how the basal ganglia and the cerebellum dynamically encode temporal information in conjunction with the premotor areas.

## Acknowledgments

We are very grateful for the valuable comments that Vani Rajendran provided to our manuscript. We also thank Raul Paulín for his technical assistance. This work was supported by Consejo Nacional de Ciencia y Tecnología (CONACYT) Grant CONACYT: A1-S-8430, UNAM-DGAPA-PAPIIT IN201721, and SECITI 2342.

